# SHIP1 partitions between cortical oscillations and membrane puncta to regulate neutrophil chemotaxis

**DOI:** 10.64898/2026.05.25.727507

**Authors:** Althea Amaris, Grace L. Waddell, Asa J. Noteboom, Sophia Doerr, Emma E. Drew, Sean R. Collins, Scott D. Hansen

## Abstract

As frontline defenders in the innate immune system, neutrophils traffic to sites of infection by detecting chemical gradients, polarizing, and migrating towards foreign pathogens. This requires communication between small GTPases, phosphatidylinositol phosphate (PIP) lipids, and the actin cytoskeleton to generate forces that drive leading-edge membrane protrusions. Central to the coordination of these signaling events is the hematopoietic-cell-specific 5-inositol lipid phosphatase SHIP1 (or INPP5D), which shapes PIP lipid domains by dephosphorylating phosphatidylinositol-(3,4,5)-trisphosphate (PIP_3_) to generate PI(3,4)P_2_. To decipher mechanisms controlling SHIP1 membrane localization during neutrophil chemotaxis, we combined genetic manipulation of SHIP1 with live cell Total Internal Reflection Fluorescence (TIRF) microscopy. Our results reveal two modes of SHIP1 localization: (1) cortical oscillations regulated by the membrane curvature sensing protein FBP17, and (2) dynamic membrane puncta organized by the adaptor protein Nck1. Localization to both membrane structures requires the proline-rich C-terminal domain (CTD) of SHIP1. Neutrophil-differentiated HL60 cells solely expressing a cytoplasmic localized SHIP1 CTD deletion mutant exhibit chemotaxis defects that phenocopy complete SHIP1 loss of function. Overall, this work reveals a dual mechanism by which SHIP1 integrates signals from membrane curvature sensing and adaptor proteins to control neutrophil cell migration.

## INTRODUCTION

The innate immune system relies on leukocytes to recognize and destroy invading threats, such as viruses, bacteria, and even cancer cells. Central for this role, these cells follow environmental signals to migrate directionally (chemotaxis) to target locations. The ability of leukocytes to transiently alter the concentration and spatial distribution of peripheral membrane proteins is critical for chemotaxis, by allowing the cell to orient, polarize, and migrate. This cellular response requires intimate communication between cell surface receptors, small GTPases, and phosphatidylinositol phosphate (PIP) lipids at the plasma membrane. Deciphering the spatiotemporal organization of these signaling networks is critical for understanding how cell polarity is established, maintained, and modulated during immune cell signaling.

In neutrophils, activation of the formyl peptide receptor by chemoattractant peptides such as N-formyl-methionine-leucine-phenylalanine (fMLF) results in the activation of phosphoinositide 3-kinase (PI3K) by GβGγ and Rho family GTPases (Rickert et al. 2000; Weiner et al. 2002; Li et al. 2003). Once active, PI3K rapidly produces phosphatidylinositol-(3,4,5)-trisphosphate (PI(3,4,5)P_3_) at the leading edge resulting in polarized F-actin polymerization, pseudopod extension, and cell migration in the direction of the chemoattractant gradient (Servant 2000; Yoo et al. 2010; Pollard and Borisy 2003; Stephens et al. 1991). However, tight spatial and temporal regulation of PI(3,4,5)P_3_ lipids is critical to focus protrusion activity for efficient migration.

Two lipid modifying enzymes – phosphatase and tensin homolog (PTEN) and Src homology 2 domain-containing inositol 5-phosphatase 1 (SHIP1) – limit the concentration of PI(3,4,5)P_3_ at the plasma membrane (Maehama and Dixon 1998; Damen et al. 1996). PTEN converts PI(3,4,5)P_3_ to PI(4,5)P_2_ and localizes to the rear of migrating *Dictyostelium* amoeba (Funamoto et al. 2002). Amoeba lacking PTEN are unable to polarize and directionally migrate (Iijima and Devreotes 2002). It is, therefore, surprising that neutrophils lacking PTEN display only subtle changes in cell migration speed (Nishio et al. 2007; Collins et al. 2015). SHIP1 dephosphorylates PI(3,4,5) P_3_ to generate PI(3,4)P_2_ (Damen et al. 1996; Pauls and Marshall 2017). In contrast to PTEN, neutrophils lacking SHIP1 display an accumulation of PI(3,4,5)P_3_, atypical F-actin polymerization across the membrane, and defects in adhesion-based cell migration (Nishio et al. 2007; Lam et al. 2012; Collins et al. 2015; Mondal et al. 2012). Neutrophils lacking SHIP1 display a large, flattened, and hyperprotrusive cell polarity phenotype and increased adhesion to extracellular matrix proteins (Nishio et al. 2007). Studies using neutrophil cell lysate show that 5-phosphatases contribute the largest amount (>90%) of the total PI(3,4,5)P_3_ phosphatase activity (Stephens et al. 1991), consistent with SHIP1 representing the major pathway for PI(3,4,5)P_3_ degradation in neutrophils. Therefore, it is critical to understand how SHIP1 is regulated, activated, and recruited to the plasma membrane to fully understand the mechanisms of cell polarity and migration in leukocytes.

In mast cells, SHIP1 and the F-BAR (Bin/ Amphiphysin/Rvs) domain-containing protein FBP17 (or FNBP1) display cortical oscillations that are regulated by phosphoinositide abundance and turnover (D. Xiong et al. 2016; M. Wu et al. 2013). The product of SHIP1 phosphatase activity, PI(3,4)P_2_, is proposed to function as a molecular timer that regulates the frequency and amplitude of cortical oscillations (D. Xiong et al. 2016). Cortical oscillations are a hallmark of excitability and involve repetitive cycles of protein recruitment, activation, and inactivation at the plasma membrane to regulate cell polarity and migration (Y. Xiong et al. 2010; Devreotes et al. 2017). Excitability has been observed in neutrophils in the form of self-organized actin waves (Weiner et al. 2007), and a critically important GTPase for cell navigation, Cdc42, exhibits local excitability in neutrophils (H. W. Yang et al. 2016). Research in *Dictyostelium*, fibroblasts, and immune cells has revealed that regulators of the actin cytoskeleton (Weiner et al. 2007), PIP lipids (Arai et al. 2010), small GTPases (M. Wu et al. 2013; H. W. Yang et al. 2016), and membrane curvature sensing proteins (Z. Wu et al. 2018; D. Xiong et al. 2016) also display features of excitability, such as all-or-nothing responsiveness and wave propagation (Devreotes et al. 2017; Y. Yang and Wu 2018). The above results suggest that these molecules are part of a conserved excitable signaling network.

SHIP1 is a multidomain protein that contains several proposed lipid binding domains and protein interaction motifs thought to regulate membrane localization and catalytic activity. We previously showed that the central catalytic core of SHIP1, which is flanked by the pleckstrin-homology related (PH-R) and C2 domains, exhibits very transient plasma membrane dwell times (Waddell et al. 2023). In the context of immune cell signaling, protein interactions mediated by the N-terminal Src homology 2 domain (SH2) and C-terminal domain (CTD) are thought to regulate SHIP1 localization. In B lymphocytes, plasma membrane localization of SHIP1 is mediated by the N-terminal SH2 domain, which interacts with immunoreceptor tyrosine-based inhibitory motifs (ITIMs) and immunoreceptor tyrosine-based activating motifs (ITAMs) (Ono et al. 1996, 1997; Nakamura et al. 2002; Isnardi et al. 2006; Mukherjee et al. 2012). The CTD contains two NPXY motifs that can facilitate interactions with phosphotyrosine-binding (PTB) domains (Manno et al. 2016; Lamkin et al. 1997), as well as proline-rich motifs that can bind SH3 domain containing proteins (Harmer and DeFranco 1999). In the context of neutrophil cell migration, it’s unclear which SHIP1 domains or motifs are required for robust plasma membrane localization.

Using neutrophil-differentiated HL-60 (dHL-60) cells as a model system, we performed a molecular dissection of SHIP1 to determine which domains or motifs control plasma membrane localization. When overexpressed in neutrophils, SHIP1 localizes to the leading edge of actin-based membrane protrusions and into cortical oscillations that sweep across the plasma membrane with a period length of 15-30 seconds/ cycle. These oscillations are spatially and temporally coupled to Cdc42(GTP) and FBP17 in the excitable signaling network. However, when SHIP1 is expressed at endogenous levels, cortical oscillations are rarely observed. Rather, SHIP1 localizes predominantly to membrane puncta along with the adaptor protein, Nck1. Puncta dynamics are sensitive to inhibitors that block integrin-mediated adhesion to substrate, suggesting that SHIP1 localizes to a macromolecular assembly that regulates adhesion receptor function. Genetic perturbations reveal that the expression levels of FBP17 and Nck1 regulate the extent to which SHIP1 partitions between cortical membrane oscillations and membrane puncta. Mutational analysis of SHIP1 indicates the C-terminal tail of SHIP1 is required for both localization patterns. Deletion of the SHIP1 C-terminal domain (CTD) containing proline-rich motifs causes defects in neutrophil chemotaxis that phenocopy SHIP1 complete loss of function. Overall, these results reveal a critical role for the CTD of SHIP1 in localizing the protein to different membrane structures based predominantly on interactions with SH3 domain containing proteins. Genetic perturbations that shift the steady-state distribution of SHIP1 interaction partners can modulate SHIP1 localization to regulate neutrophil cell migration.

## RESULTS

### SHIP1 plasma membrane localization visualized in differentiated neutrophil-like cells

A major limitation in our understanding of the role SHIP1 plays in neutrophil chemotaxis is that the mechanisms controlling its plasma membrane localization during neutrophil cell polarity and migration remain unclear. Previous research imaged SHIP1 localization patterns in migrating neutrophils in zebrafish larvae (Lam et al. 2012), however this study lacked the spatial resolution needed to define the mechanisms that control SHIP1 membrane recruitment and dynamics. Similarly, previous studies of SHIP1 localization in B cells and mast cells were restricted to chemically fixed or non-motile cells (Pauls et al. 2020; D. Xiong et al. 2016). To decipher the mechanisms that regulate SHIP1 localization during chemotaxis, we established methods to visualize SHIP1 in dHL-60 cells using Total Internal Reflection Fluorescence (TIRF) microscopy. In the presence of fibronectin coated glass surfaces, dHL-60 cells adhere through integrin receptors. When these cells were uniformly stimulated with fMLF, exogenously expressed mNeonGreen-SHIP1 (mNG-SHIP1) localized to the leading-edge of actin-based membrane protrusions (**Fig. 1A**). This mNG-SHIP1 localization pattern was also observed in randomly migrating cells were imaged using an under-agarose preparation in the absence of fibronectin.

**Figure 1.**
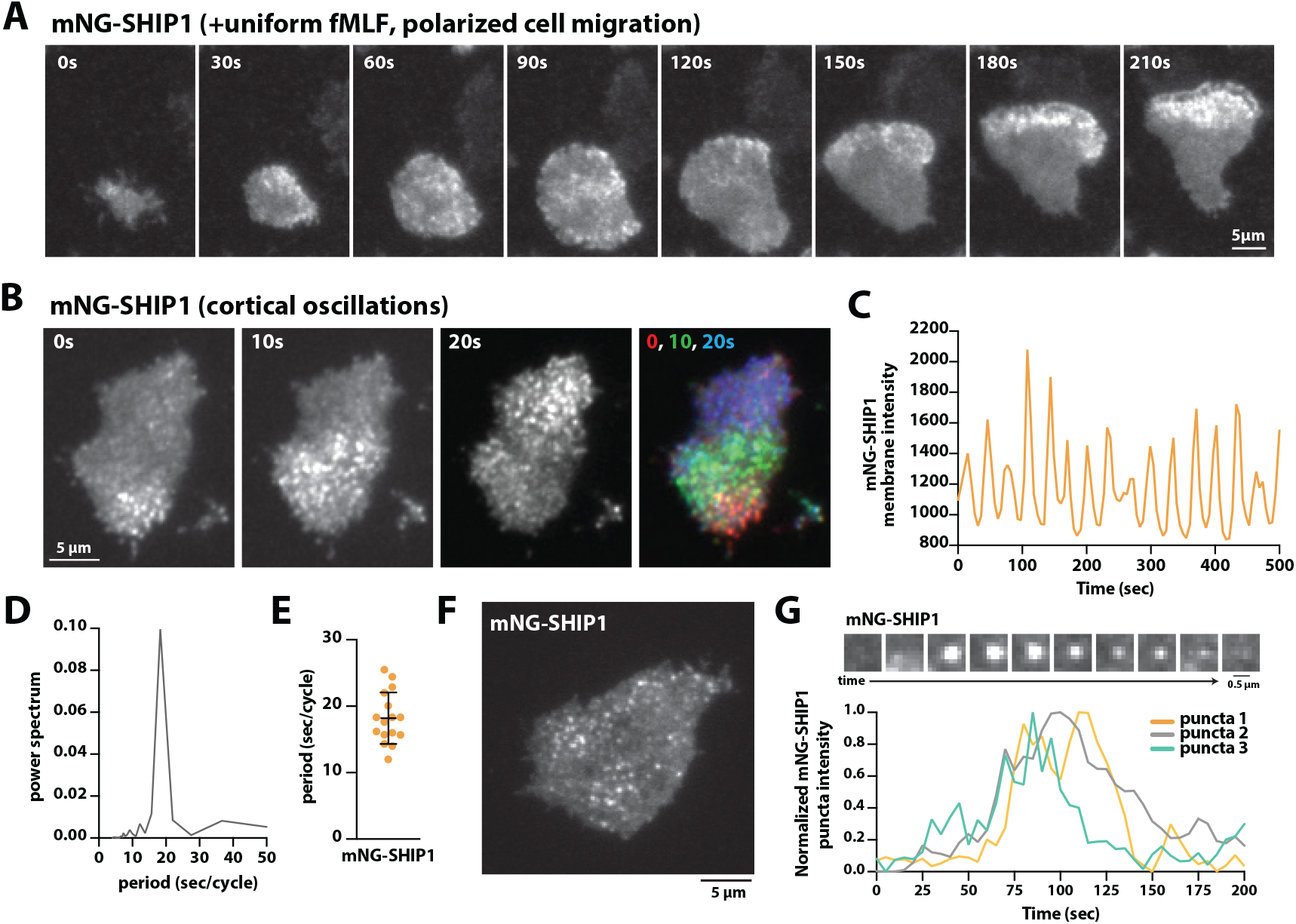
SHIP1 localizes to actin-based membrane protrusions, cortical oscillations, and puncta in differentiated neutrophil-like cells. **(A)** Stimulation of dHL-60 cells with a uniform concentration of 20 nM fMLF promotes polarized leading edge membrane localization of mNG-SHIP1. Cells were adhered to fibronectin coated coverslips. Visualized by TIRF-M. **(B)** mNG-SHIP1(1-1189aa) localizes to cortical membrane oscillations in dHL-60 cells. Overlay of multiple frames shows wave movement. **(C)** Graph showing time dependent fluctuations (i.e. cortical oscillations) in mNG-SHIP1 plasma membrane localization. **(D)** Fourier transform of data in **(C)** reveals characteristic frequency (or period length) of mNG-SHIP1 cortical oscillations. **(E)** Quantification of mNG-SHIP1 oscillation frequency (n = 16 cells, 18.2 ± 3.9 sec). **(F)** mNG-SHIP1 displays punctate localization at the plasma membrane. **(G)** Representative mNG-SHIP1 membrane puncta rise and fall in fluorescent intensity.

Neutrophil-like cells that ectopically express mNG-SHIP1 predominantly displayed cortical oscillations at the plasma membrane (**Fig. 1C** and **D**). Fourier transform analysis of the mNG-SHIP1 membrane intensity profile revealed a period length of 18 seconds/cycle (**Fig. 1E**). This behavior was similar to the localization pattern previously observed for SHIP1 exogenously expressed in mast cells (D. Xiong et al. 2016). Cortical oscillations of mNG-SHIP1 were never observed in undifferentiated HL-60 (uHL-60) cells, indicating that the gene expression profile associated with the differentiated neutrophil-like state is critical for establishing these membrane localization dynamics.

After culturing neutrophils that ectopically express mNG-SHIP1 for more than a week, we observed a gradual reduction in the cellular concentration of mNG-SHIP1 in our live cell imaging experiments. Similar results were observed using either the SFFV or ubiquitin promoters. The gradual downregulation of exogenously expressed mNG-SHIP1 revealed small diffraction-limited membrane puncta (**Fig. 1F**). These mNG-SHIP1 puncta displayed characteristic plasma membrane intensity dynamics which included a rise and fall in membrane intensity (**Fig. 1G**), reminiscent of the dynamics observed for a variety of endocytic proteins (Kaksonen et al. 2003, 2005; Taylor et al. 2011). Using a range of mNG-SHIP1 expression conditions, we aimed to determine which SHIP1 domains were required for localization to actin-based membrane protrusions, cortical oscillations, and membrane puncta.

### The C-terminal domain is necessary and sufficient for robust SHIP1 membrane localization

SHIP1 is a multidomain protein that has several proposed lipid and protein interaction domains (**Fig. 2A**), which could mediate localization to cortical oscillations and to the leading edge. In our previous biochemical analysis of SHIP1, we found that the protein displays a very weak affinity for phosphatidylserine (PS) and PI(3,4,5)P_3_ lipids presented on a reconstituted supported lipid bilayer (Waddell et al. 2023). To explore the role of these interactions in living cells, we conducted single molecule dwell time analysis of mEos-SHIP1 constructs. A construct containing the central phosphatase domain with flanking PH and C2 domains revealed transient interaction with the plasma membrane. In contrast, the dwell time for full-length mEos-SHIP1 was approximately 20-fold longer compared to mEos-SHIP1 (PH-PP-C2) (**Fig. 2B-2C**). This suggests that protein interactions are critical for regulating plasma membrane localization of SHIP1 in vivo. To determine the domain/motif(s) required for SHIP1 localization in dHL-60 cells, we performed a molecular dissection. Although the SH2 domain was previously shown to be required for SHIP1 plasma membrane localization in B-cells (Mukherjee et al. 2012), we found that this domain was dispensable for SHIP1 localization to both actin-based membrane protrusions and cortical oscillations (**Fig. 2D-2F**). By contrast, mNG-SHIP1 mutants lacking the C-terminus were uniformly localized to the plasma membrane in a manner that was indistinguishable from mNG-SHIP1 (PH-PP-C2) (**Fig. 2D**).

**Figure 2.**
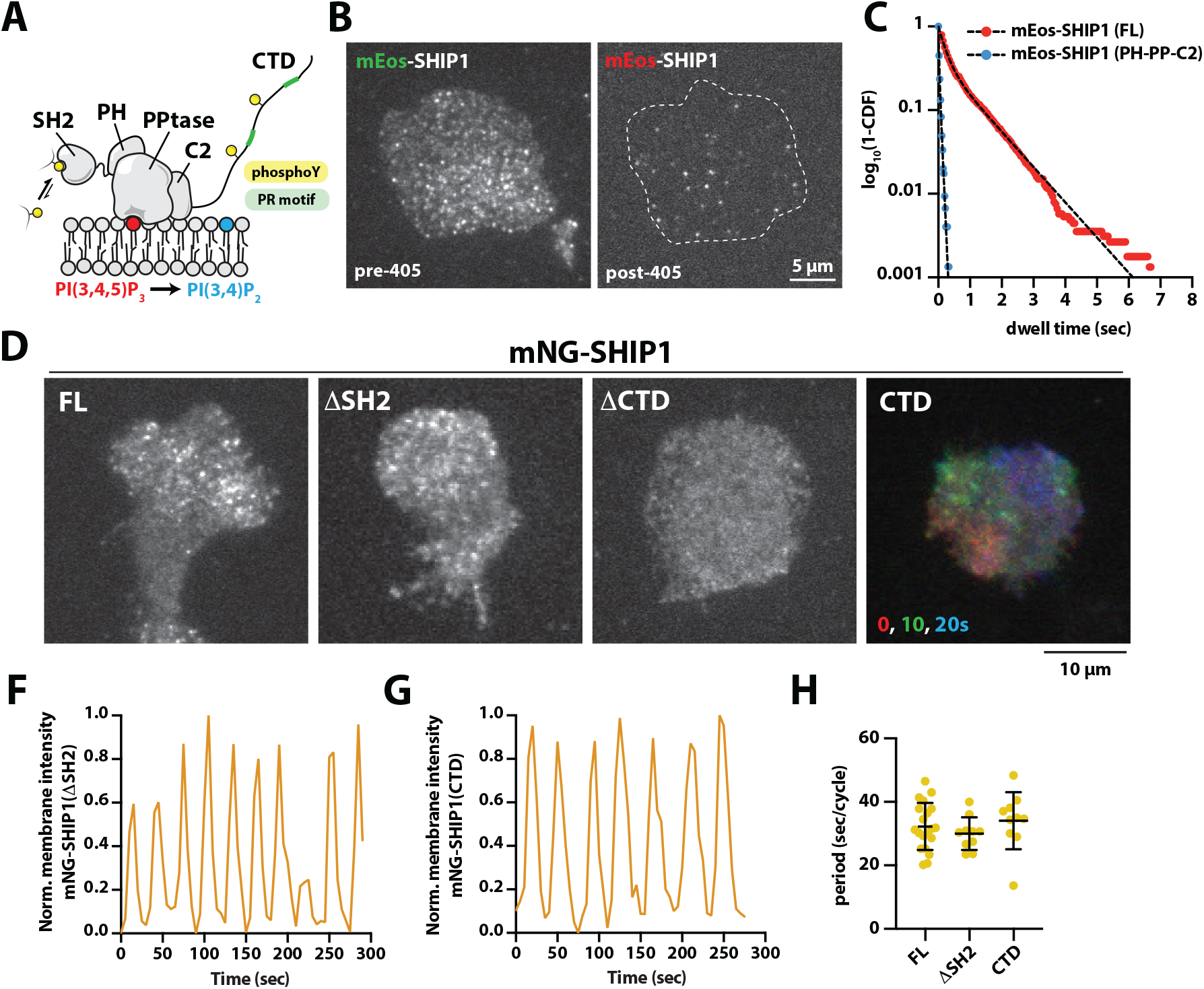
The C-terminal domain is necessary and sufficient for robust SHIP1 membrane localization. **(A)** Proposed structural organization of SHIP1 (1-1189aa). **(B)** Representative TIRF-M images showing the localization of mEos-SHIP1 (FL, 1-1189aa) before (left) and after (right) photoconversion with 405 nm light. **(C)** Single molecule dwell time distribution of mEos-SHIP1(FL) localized to the plasma membrane in dHL-60 cells. Data plotted as 1-log_10_ cumulative distribution frequency (CDF) and fit to single and double exponential curves yielding the following dwell times: τ_1_=0.03 sec (n = 1543 mEos-SHIP1 PH-PP-C2 molecules); τ_1_=0.23 sec, τ_2_=1.03 sec, α=0.62 (n = 2260 mEos-SHIP1 FL molecules). **(D)** Representative TIRF-M images showing the localization of mNG-SHIP1(1-1189aa), ΔSH2(102-1189aa), ΔCTD(1-1078aa), and CTD(1079-1189aa) in dHL-60 cells. Overlay of multiple frames shows wave movement mNG-SHIP1(CTD,1079-1189aa). **(F-G)** Graph showing cortical oscillations of **(F)** mNG-SHIP1(ΔSH2,102-1189aa) and **(G)** mNG-SHIP1(CTD,1079-1189aa) at the plasma membrane. **(H)** Quantification of mNG-SHIP1 oscillation frequency.

The C-terminus of SHIP1 is a predicted random coil, but it contains several predicted phosphotyrosine sites (i.e. NPXY motif) that could facilitate protein-protein interactions to regulate SHIP1 membrane localization. Although post-translational modifications of SHIP1 have not been mapped in human neutrophils, the C-terminus of SHIP1 is reportedly tyrosine phosphorylated in other hematopoietic cells (Lamkin et al. 1997; Manno et al. 2016). Mutating one reported phosphotyrosine site, Y1022A, revealed no change in SHIP1 membrane localization. In addition to the predicted phosphotyrosine sites, the C-terminus of the SHIP1 contains several poly-proline motifs that could interact with Src homology 3 (SH3) domain containing proteins (Teyra et al. 2017). Mutating the previously characterized and uncharacterized proline-rich motifs did not disrupt SHIP1 localization into cortical oscillations. To identify the minimal motif required for SHIP1’s polarized membrane localization pattern, we created a series of C-terminal truncations. This revealed that the last 110 amino acids of the SHIP1 C-terminus are required for localization to actin-based membrane protrusions, cortical oscillations, and membrane puncta. A SHIP1 mutant lacking this region localized uniformly in the absence and presence of chemoattractant (**Fig. 2D**). To test whether the 110aa C-terminal fragment was sufficient for localization to cortical membrane oscillation, we expressed this fragment as a mNG fusion. Although the signal-to-noise ratio was reduced, mNG-SHIP1 (1079-1189aa) displayed cortical membrane oscillations with a frequency similar to that of full length SHIP1 (**Fig. 2G-2H**). However, membrane puncta localization of mNG-SHIP1 (1079-1189aa) was not observed (**Fig. 2D**).

### The C-terminus is required for SHIP1’s role in neutrophil chemotaxis

To determine the functional significance of the C-terminal domain in controlling SHIP1 plasma localization in neutrophils, we established CRISPR interference (CRISPRi) to silence expression of endogenous SHIP1 in dHL-60 cells using multiple guide RNAs (gRNAs) (**Fig. 3A**). Our set of gRNAs also included a sequence targeting the SHIP2 locus to limit genetic compensation resulting from the loss of SHIP1. Western blot and RT-qPCR analysis indicated a 95% and 50% knockdown efficiency of SHIP1 and SHIP2, respectively (**Fig. 3A**). Consistent with these cells having an elevated level of PI(3,4,5) P_3_, these CRISPRi cells exhibited a hyperprotrusive phenotype as previously reported (Nishio et al. 2007), characterized by the formation of multiple pseudo-leading edges and a corresponding drop in overall cell solidity (a shape parameter) as compared to wild-type (**Fig. 3B-3C**). Mutant cells visualized under agarose also displayed a larger membrane surface area and a less spherical morphology compared to the control cells, exhibiting up to a four-fold increase in 2D spread area (**Fig. 3B-3C**). Using this cell line, we discovered that prolonged knockdown (KD) by CRISPRi could not be rescued through exogenous expression of mNG-SHIP1. Although cells could be infected with lentivirus to express mNG-SHIP1 after stable SHIP1 KD by CRISPRi, these cells continued exhibiting cell migration defects. This highlights the challenges associated with generating a complete SHIP1 knockout cell line for the purpose of mutant rescue experiments. As previously reported, cells with persistently high levels of PI(3,4,5) P_3_ and hyperactivated Akt undergo epigenetic changes that alter global gene expression (Kiselev et al. 2015) and proteome interactions (Swaney et al. 2021).

**Figure 3.**
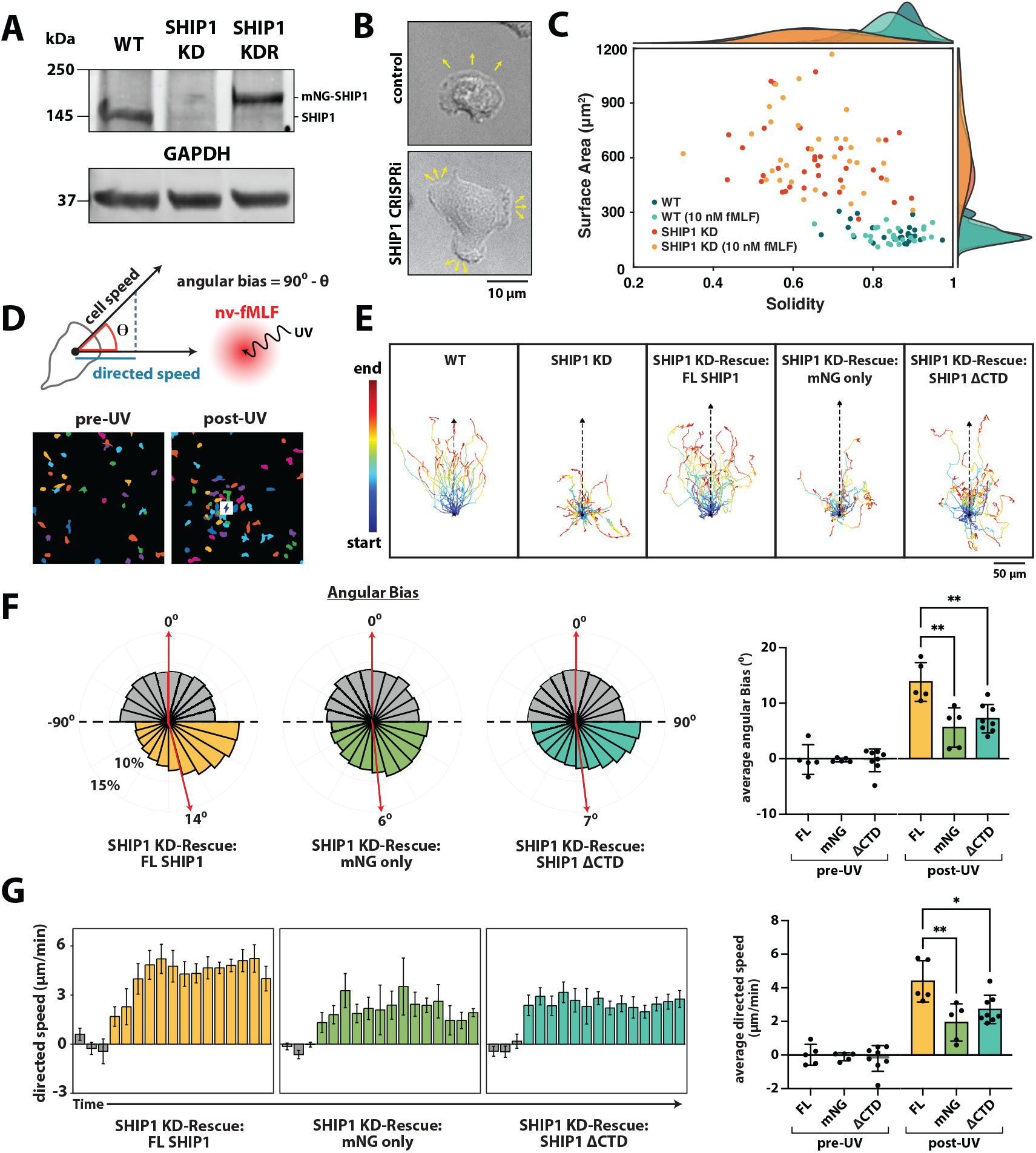
Neutrophils expressing mNG-SHIP1ΔCTD) exhibit chemotaxis defects. **(A)** Western blot showing SHIP1 protein levels in wild type dHL-60 cells, CRISPRi knockdown targeting SHIP1, and subsequent rescue with mNG-SHIP1. **(B)** Representative DIC images of wildtype and SHIP1 CRISPRi cells. Arrows indicate the direction of leading-edge protrusions. **(C)** Morphological quantification showing difference in cell surface area and solidity in dHL-60 with SHIP1 CRISPRi. **(D)** nv-fMLF is uncagged with UV light to generate a local chemoattractant concentration gradient of fMLF. Diagram showing the experimental approach for quantifying light-induced chemotaxis. Cell speed and directed speed are quantified, respectively, as the magnitude of the velocity vector and the velocity component projected along the cell-chemoattractant axis. Angular bias scales from 0 to 90; 0 represents random movement and 90 represents the ideal trajectory toward the chemoattractant source. (Below) Cell masks, segmented using Cellpose, depicting cell positions relative to the chemoattractant source pre- and post-UV-uncaging. **(E)** Representative single cell trajectories (n = 25-30 cells) showing neutrophil chemotaxis in fMLF gradient. The black triangle represents median chemoattractant distance relative to initial cell position. **(F)** Mirrored polar histograms showing angular bias pre- (gray) and post-uncaging (colored). Red line is population mean angular bias (n = 445 (KDR:FL), 88 (KDR:mNG), 478 (KDR:ΔCTD) total cells, 5-8 independent experiments). **(G)** Time-course of dHL-60 directed speed pre- (gray) and post-uncaging (colored). Error bars denote mean (n = 445 (KDR:FL), 88 (KDR:mNG), 478 (KDR:ΔCTD) total cells, 5-8 independent experiments). Bins equal 90 seconds. **(F, G)** Average angular bias **(F)** and directed speed **(G)** pre- (gray) and post-uncaging (colored). **(F, G)** Circular markers represent independent experiments (n = 10 - 140 cells per experiment). Error bars equal mean +/-SD. p* < 0.05, p** <0.01 (one-way ANOVA with Tukey’s post hoc test).

To overcome the limitations of our initial approach, we generated a CRISPRi construct to simultaneously silence SHIP1 expression and rescue with either wild-type or mutant mNG-SHIP1. Using this approach, we were able to match endogenous SHIP1 expression levels (**Fig. 3A**) and overcome the major defects described above. Consistent with initial localization studies of overexpressed mNG-SHIP1, the C-terminus was required for mNG-SHIP1 localization of plasma membrane puncta (**Fig. 3C**). Notably, the localization of mNG-SHIP1 expressed at levels comparable to the endogenous protein did not exhibit strong cortical membrane oscillations (**Fig. 3C**). This indicates that SHIP1 is primarily localized to membrane puncta in dHL-60 cells.

To determine the functional significance of CTD-mediated localization of SHIP1 in regulating neutrophil chemotaxis, we used photocaged *N*-nitroveratryl fMLF (nv-fMLF) in under-agarose chemotaxis assays (Bell et al. 2018; Collins et al. 2015). Following a pulse of UV light in a spatially defined location, the chemoattractant is released, generating a localized chemoattractant gradient (**Fig. 3D**). This stimulates neutrophil polarity and migration towards the uncaging site (**Fig. 3E**). While wild-type cells rapidly oriented and migrated towards the uncaging site, SHIP1 KD cells exhibited chemotactic defects. These cells frequently failed to orient properly towards the chemoattractant source, and they rarely reached the site of uncaging. Crucially, CRISPRi cells rescued with full-length (FL) SHIP1 completely rescued chemotaxis, whereas CRISPRi cells rescued with only mNG continued to exhibit the same chemotaxis defects as the knockdown cells (**Fig. 3E-3G**). To evaluate these defects, we quantified both the angular bias (the tendency of a cell to orient towards the chemoattractant source) and the directed speed (the component of velocity in the direction of the gradient) (**Fig. 3F-3G**). The rose plot distributions of single-cell trajectories revealed that prior to chemoattractant uncaging, all cell lines exhibited a symmetrical, unpolarized distribution with a mean angular bias of approximately 0°, consistent with random migration (**Fig. 3F**). Following uncaging, the angular bias distribution of the FL mNG-SHIP1 rescue cells shifted asymmetrically towards the 90° axis (which represents the gradient center), averaging 14° (**Fig. 3F**). In contrast, rescue cells expressing only mNG or the mNG-SHIP1(ΔCTD) mutant lacked this directional fidelity. While both rescues retained the ability to sense the gradient, as seen by a very slight right lean in their angular bias distributions compared to their pre-uncaging baselines, their trajectories were less directional (**Fig. 3F**). Compared to each other, the mNG and mNG-SHIP1(ΔCTD) rescues yielded statistically insignificant post-uncaging average angular biases of 6° and 7° respectively, showing that both constructs fail to restore wild-type orientation (**Fig. 3F**).

This loss of direction-sensing was directly mirrored in the directed migration speeds. While the directed speed of FL SHIP1 rescue cells rapidly increased to an average of 4.39 µm/min, the directed speeds of the mNG and mNG-SHIP1(ΔCTD) mutant cells plateaued earlier (**Fig. 3G**), yielding significantly lower average directed speeds of 1.94 and 2.72 µm/min respectively (**Fig. 3G**). Importantly, the chemotaxis defects were not caused by the mutant cells being unable to move. Quantification of total, non-directed migration speed revealed that post-uncaging, all three rescue cell lines exhibited a chemokinetic response that increased their own overall speeds. However, the post-uncaging average migration speeds of the mNG and mNG-SHIP1(ΔCTD) mutant cells were not statistically different from that of the FL mNG-SHIP1 rescue. Together, these metrics confirm that while cells lacking the SHIP1 C-terminus remain motile, they exhibit a loss of function and fail to rescue the wild-type chemotaxis phenotype. The C-terminus of SHIP1 is therefore required for proper neutrophil chemotaxis.

### SHIP1 localization is modulated by membrane curvature sensing proteins

Previous research in mast cells revealed that FBP17 localizes to the plasma membrane and regulates cortical oscillations downstream of the active Cdc42 GTPase, PIP lipids, and membrane topographical features (M. Wu et al. 2013; D. Xiong et al. 2016; Z. Wu et al. 2018). Based on coimmunoprecipitation, SHIP1 was shown to interact with the SH3 domain derived from FBP17 in cell lysate (D. Xiong et al. 2016). To determine whether SHIP1 localizes to cortical oscillations similarly in neutrophil-like cells, we visualized the dynamics of active Cdc42 in dHL-60 cells. Previous research examining the role of Ras/Rho-GTPases in neutrophil cell polarity and migration found that a Cdc42 FRET biosensor exhibited cortical membrane oscillations (H. W. Yang et al. 2016). Expression of an iRFP tagged CRIB (Cdc42/Rac Interactive Binding) domain from N-WASP with specificity for Cdc42(GTP) exhibited similar cortical membrane oscillations (**Fig. 4A-4B**). Co-expression of mNG-SHIP1 and iRFP-CRIB revealed cortical membrane oscillations that were spatially and temporally coupled (**Fig. 4B**) with mNG-SHIP1 localization lagging slightly behind the Cdc42(GTP) sensor. We also observed a similar degree of spatiotemporal coupling when GFP-FBP17 and iRFP-CRIB were co-expressed. Treatment of neutrophils with an allosteric inhibitor that blocks GEF dependent activation of Cdc42 perturbed the localization of both SHIP1 and FBP17 into cortical oscillations.

**Figure 4.**
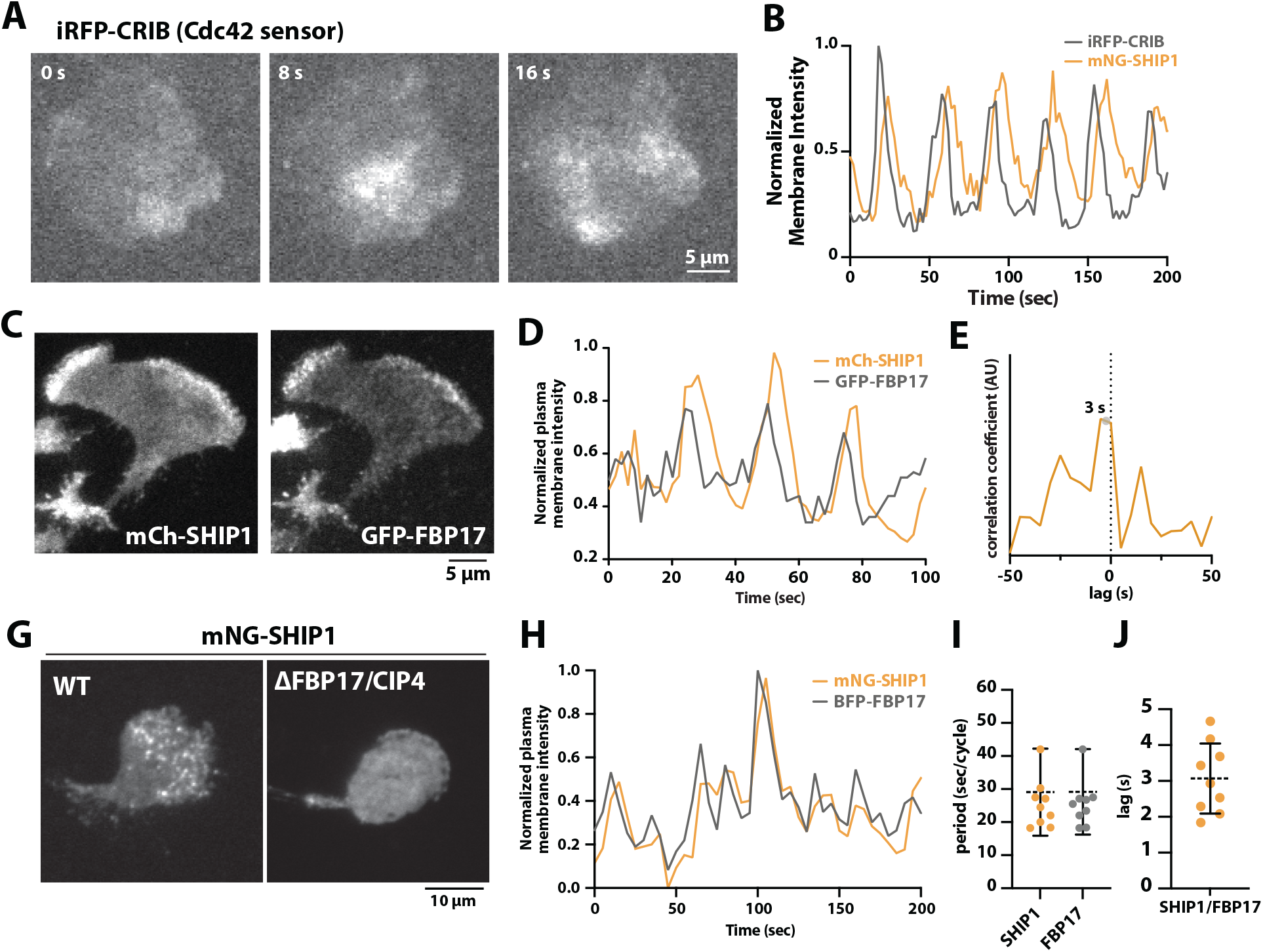
FBP17 regulates SHIP1 localization to membrane puncta and cortical oscillations. **(A)** Cortical oscillation of iRFP-CRIB (Cdc42(GTP) sensor) visualized in dHL-60 cells. **(B)** SHIP1 and Cdc42(GTP) exhibit cortical membrane oscillations. Normalized iRFP-CRIB and mNG-SHIP1 membrane intensity fluctuations visualized in dHL-60. **(C)** mCherry-SHIP1 and GFP-FBP17 colocalize to actin-based membrane protrusions when overexpressed in dHL-60. **(D)** mCherry-SHIP1 and GFP-FBP17 display synchronized cortical oscillations when overexpressed in dHL-60 cells. **(E)** Representative cross-correlation plot derived from dHL-60 overexpressing mCherry-SHIP1 and EGFP-FBP17. Time indicates lag in SHIP1 localization relative to FBP17. **(G)** mNG-SHIP1 localizes predominantly to membrane puncta when expressed at low levels in dHL-60 cells. mNG-SHIP1 membrane puncta localization is perturbed in FBP17Δ/CIP4Δ dHL-60 cells. **(H)** mNG-SHIP1 localization can be shifted from predominantly membrane puncta to cortical membrane oscillation by overexpressing BFP-FBP17 in dHL-60 cell. Membrane intensity fluctuations of mNG-SHIP1 and BFP-FBP17 overexpressed in dHL-60 cell. **(I)** Quantification of period length for mNG-SHIP1 (n = 10 cells, 29.1 ± 13.2 sec). and BFP-FBP17 (n = 10 cells, 29.2 ± 12.9 sec). **(J)** Quantification of lag time between the arrival of GFP-FBP17 and mCherry-SHIP1 (n = 9 cells, 3.1±1 sec).

The primary sequence and domain architecture of SHIP1 suggests that it does not directly interact with Cdc42. However, FBP17 contains a Rho-binding extended motif (REM) that was previously shown to bind Cdc42(GTPγS) using purified proteins (M. Wu et al. 2013). When we ectopically co-expressed GFP-FBP17 and mCherry-SHIP1 in dHL-60 cells, the two proteins colocalized to the leading edge of actin-based membrane protrusions (**Fig. 4C**). Cortical oscillations were spatially and temporally synchronized with mCherry-SHIP1 lagging slightly behind peak GFP-FBP17 by about 3 seconds (**Fig. 4D-4E**). Notably, the co-expression of mCherry-SHIP1 and GFP-FBP17 synergistically exaggerated the cortical oscillations. The localization of both proteins was enhanced compared to cells expressing the individual fluorescently tagged proteins. Under overexpression conditions, cortical oscillations of SHIP1 are prominent, while membrane puncta are challenging to resolve.

To determine the epistatic relationship between SHIP1 and FBP17, we visualized the localization of mNG-SHIP1 in differentiated HL-60 (dHL-60) neutrophil-like cells containing a genetic deletion of FBP17 and the functionally redundant membrane curvature sensing protein denoted CIP4 (Brunetti et al. 2022). In wild type dHL-60s, we found that mNG-SHIP1 localized primarily to membrane puncta when ectopically expressed (**Fig. 4G**). In this context, cortical oscillations of mNG-SHIP1 were rarely observed. This localization pattern phenocopied the CRISPRi knockdown-rescue cell line expressing mNG-SHIP1. Visualization of mNG-SHIP1 localization in dHL-60s lacking FBP17 and CIP4, revealed a complete loss of mNG-SHIP1 membrane puncta (**Fig. 4G**). In contrast, overexpression of BFP-FBP17 in the mNG-SHIP1 CRISPRi knockdown-rescue cell line resulted in the redistribution of mNG-SHIP1 from predominantly membrane puncta to cortical membrane oscillations (**Fig. 4H-4J**). These results indicate that SHIP1 predominantly localizes to membrane puncta. However, overexpression of the membrane curvature sensing protein, FBP17, can shift the steady-state distribution of molecular interactions to promote SHIP1 localization to cortical membrane oscillations.

### Relationship between SHIP1 membrane puncta and endocytosis

The dynamics of mNG-SHIP1 punctate membrane localization is reminiscent of proteins that regulate endocytosis. To determine whether SHIP1 localizes to sites of endocytosis, we compared its spatial distribution to canonical endocytic markers. First, we co-expressed mNG-SHIP1 with low levels of mCherry-clathrin, the master regulator of receptor-mediated endocytosis. Consistent with previous observations of endogenous clathrin localization in dHL-60s (Brunetti et al. 2022), mCherry-clathrin light chain was highly enriched at the trailing edge of randomly migrating dHL-60 cells (**Fig. 5A**). By contrast, mNG-SHIP1 puncta were strongly localized towards the leading edge of polarized neutrophils (**Fig. 5A**). To determine if SHIP1 colocalizes with other endocytic proteins, we next examined the clathrin-independent fast endophilin mediated endocytosis (FEME) pathway. The BAR domain containing protein, endophilin, is proposed to regulate vesicle scission by inducing membrane curvature, scaffolding protein-protein interactions, and recruiting dynamin to sites of endocytosis (Renard et al., 2015). Although the substrate and product of SHIP1/2 phosphatase activity have been reported to modulate FEME (Chan Wah Hak et al. 2018; Boucrot et al. 2015), SHIP1/2 have not been reported to localize to endocytic pits. The human genome encodes five endophilin isoforms denoted A1, A2, A3, B1, and B2. Based on RNA sequencing data, human primary neutrophils and dHL-60 cells express endophilin A2, B1, and B2 (Rincón et al. 2018). To characterize the localization and dynamics of these endocytic proteins, we generate dHL-60 cells that stably express low levels of endophilin A2, B1, or B2 tagged with mScarlet. While all three isoforms localized to dynamic membrane puncta visualized by TIRF microscopy, endophilin B1 and B2 puncta were relatively sparse and had a low signal-to-noise ratio (**Fig. 5A**). In contrast, endophilin A2 formed bright dynamic puncta that were abundantly distributed across the plasma membrane. This aligns with established literature identifying endophilin A2 as the primary regulator of FEME. Although endophilin A2 was shown to localize to the leading edge of actin-based membrane protrusion during epithelial and mesenchymal cell migration, we rarely observed endophilin A2 puncta at the front of polarized dHL-60 cells. Instead, endophilin A2 localization was observed throughout the cell body of randomly migrating cells. When co-expressed with mNG-SHIP1, co-localization with endophilin A2 was rarely observed (**Fig. 5B-5D**).

**Figure 5.**
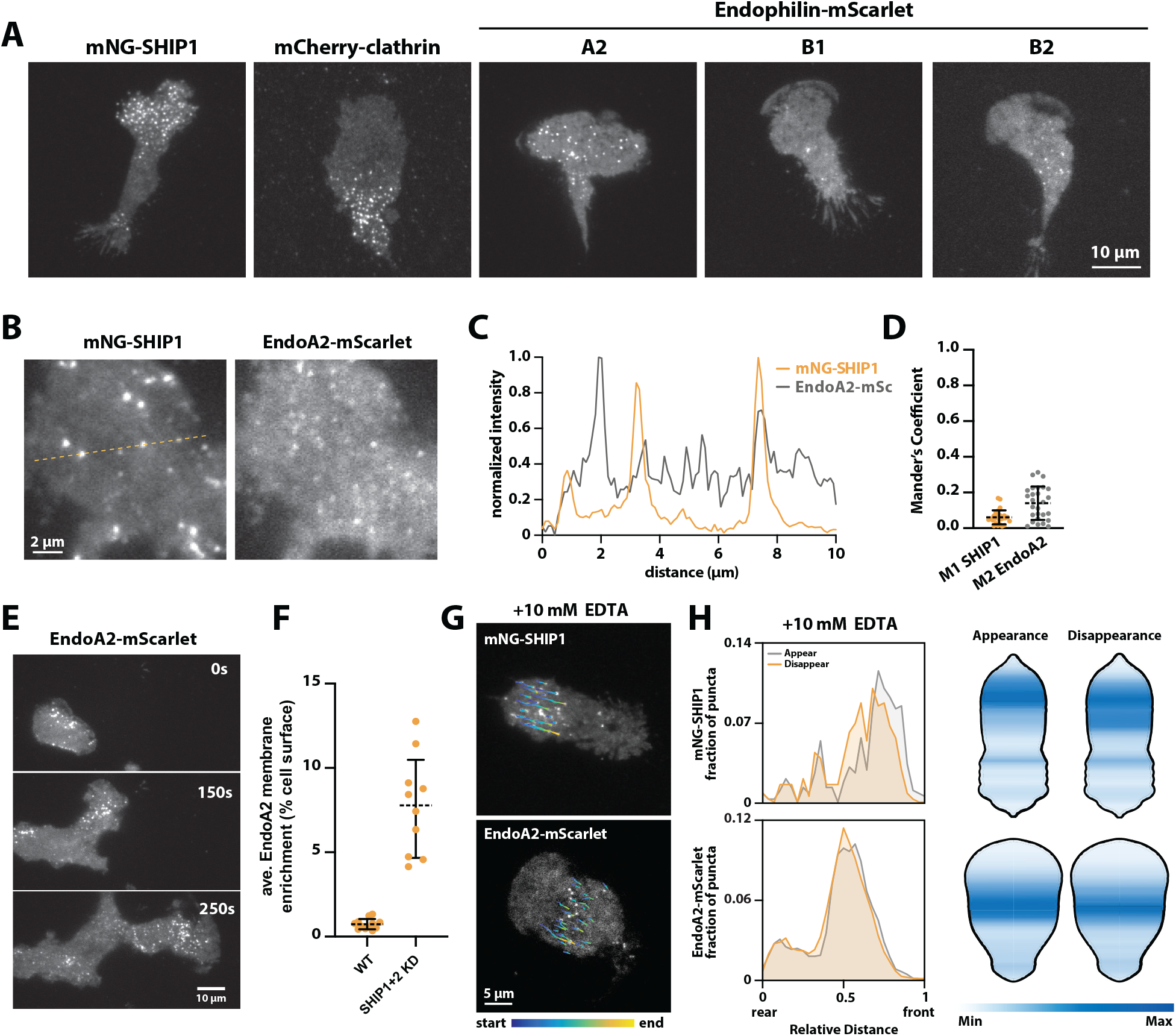
SHIP1 membrane puncta are sites of adhesion rather than endocytosis. **(A)** Representative TIRF-M images showing the punctate membrane localization of mNG-SHIP1, mCherry-clathrin, and endophilin-mScarlet isoforms A2, B1, and B2. **(B-D)** SHIP1 and endophilin A2 exhibit minimal colocalization at the plasma membrane. **(B)** Representative images showing co-expression of mNG-SHIP1 and endophilinA2-mScarlet. **(C)** Normalized fluorescence intensities corresponding to the line scan in **(B)** show that peaks of mNG-SHIP1 and endophilinA2-mScarlet are largely distinct and non-overlapping. **(D)** Mander’s coefficients quantifying the fraction of SHIP1 overlapping with endophilin A2 (M1, 0.06 ± 0.04 orange) and vice versa (M2, 0.14 ± 0.1, gray). Points equal the coefficient for a single frame (n = 9 total cells; N = 27 frames). **(E)** Representative montage of TIRF-M images showing the plasma membrane localization of endophilin-mScarlet in SHIP1 CRISPRi knockdown cells. **(F)** Quantification of exogenous endophilin-mScarlet membrane enrichment in wildtype (n = 12 cells, 0.74 ± 0.3 %) compared to SHIP1 CRIPSRi knockdown cells (n = 10 cells, 7.8 ± 3 %). **(G)** mNG-SHIP1 and endophilinA2-mScarlet puncta exhibit retrograde flow in dHL-60 cells treated with 10 mM EDTA. Color-mapped trajectories indicate track start and end (blue to yellow, maximum length). **(H)** Representative puncta appearance and disappearance probability maps relative to the rear (0) and leading edge (1) of a single cell. Line plots (left) and heatmaps (right) show mNG-SHIP1 and endoA2-mScarlet distribution throughout the cell body.Circular markers indicate individual cell averages. Bars equal mean ± SD.

Despite the lack of colocalization, SHIP1 remains mechanistically linked to endophilin activity through its regulation of phosphoinositide synthesis. Perturbing SHIP1 levels drastically altered endophilin behavior. In SHIP1 knockdown (KD) cells, where PI(3,4,5)P_3_ levels remain elevated, endophilin puncta accounted for a significantly greater fraction of the total membrane area (**Fig. 5E-5F)**, accompanied by more complex oscillatory dynamics. Interestingly, ectopic co-expression of mNG-SHIP1 with endophilinA2-mScarlet induced a similar dysregulation in endophilin resulting in an increase in membrane puncta and the onset of faint oscillatory behaviors that were entirely absent when endophilin was expressed alone. Together, this data demonstrates that while SHIP1 function is essential for initiating FEME via its lipid product. However, SHIP1 does not appear to play a direct structural role in endocytosis.

Compared to other cell types, neutrophils do not assemble long-lived focal adhesions (Yürüker and Niggli 1992). Considering the stationary localization of mNG-SHIP1 puncta relative to the migrating neutrophil, we hypothesized that SHIP1 localization could be regulated downstream of adhesion receptors. This would be consistent with SHIP1 mutant cell lines exhibiting more dramatic cell migration defects on fibronectin coated substrates (Mondal et al. 2012). To determine whether SHIP1 or endophilin A2 localize to sites of integrin-based adhesion, we disrupted cell-substrate adhesion using EDTA. By chelating divalent cations, EDTA blocks integrin-based adhesion (Zhang and Chen 2012). Without physical traction, neutrophils are unable to migrate. As a result, membrane-associated proteins tethered to the actin cytoskeleton exhibit retrograde flow towards the cell rear. Consistent with prior localization studies of clathrin-coated endocytic pits in the presence of EDTA (Brunetti et al., 2021), both endophilin A2-mScarlet and mNG-SHIP1 puncta exhibited retrograde flow (**Fig. 5G**). This is consistent with both proteins localizing to molecular assemblies that are anchored to integrin receptors and/or the actin cytoskeleton. However, when mapping the spatial probability of the puncta flux along a normalized longitudinal axis (where 0 represents the cell rear and 1 the leading edge), distinct localization patterns for SHIP1 and endophilin A2 were revealed (**Fig. 5H**). SHIP1 puncta consistently appeared near the leading edge, with appearance peaking at a relative position of ∼0.8. These puncta persisted only briefly before disappearing further back (relative position of ∼0.7) as they were dragged with the actin cytoskeleton. In contrast, endophilin A2 puncta most frequently appeared in the middle of the cell body, having both appearance and disappearance peaking at a relative position of ∼0.5. Overall, leading edge SHIP1 localization did not correlate with sites of endophilin-mediated endocytosis.

### Nck1 regulates SHIP1 localization to membrane puncta

The observation that SHIP1 localizes to both cortical oscillations and puncta at the plasma membrane suggests that distinct sets of molecular interactions drive each localization pattern. Although FBP17 is critical for the initial plasma membrane localization of SHIP1, FBP17 does not persistently localize to the plasma membrane as a punctate structure. The C-terminal domain of SHIP1 contains multiple proline-rich motifs and therefore we hypothesized that another adaptor protein regulates SHIP1 localization to plasma membrane associated puncta. Given that deletion of SHIP1’s proline-rich C-terminus both abolished SHIP1 puncta formation and resulted in a loss of chemotactic function, the formation of SHIP1 puncta is most likely driven by the protein-protein interactions that involve the C-terminal proline-rich motifs. Among the candidates, the Nck1/2 adaptor proteins have a conserved architecture consisting of one SH2 domain and three SH3 domains (Buday et al., 2002). In B-cells, Nck1/2 was also previously shown to regulate actin turnover in collaboration with SHIP1 (Pauls et al. 2020). In several immune and motility-related signaling systems, Nck1/2 proteins are proposed to link phosphotyrosine-rich protein complexes to actin remodeling machinery, including WASP/N-WASP, WIP, PAK, and the Arp2/3 actin nucleation complex (Lettau et al., 2009; Rohatgi et al., 2001; Chaki et al., 2013). Nck1/2 and SHIP1 reportedly localized to Syk-associated signaling complexes in mast cells following receptor activation (de Castro et al., 2012). Overall, the domain organization of Nck1/2 allow binding to proline-rich and tyrosine phosphorylated proteins involved in actin cytoskeletal function.

To determine whether Nck1 regulates SHIP1 localization in neutrophils, we first quantified the spatiotemporal relationship between these proteins. Live-cell imaging of dHL-60 cells co-expressing mNG-SHIP1 and mCherry-Nck1 revealed strong colocalization to puncta structures as seen by the extent of spatial overlap and Manders’ coefficients (**Fig. 6A-6B**). Compared to SHIP1, Nck1 puncta were easier to resolve and less diffusely distributed across the cell membrane. We therefore used the Nck1 signal as a reference to mask and track puncta in both channels. To quantify the spatiotemporal dynamics of the SHIP1 and Nck1 puncta, we tracked puncta based on fluorescence intensity and lifetime. Both Nck1 and SHIP1 traces followed a similar pattern: (1) puncta appearance, (2) increase to peak membrane intensity, and (3) decrease in intensity before disappearance (**Fig. 6C**). Like the time dependent change in intensity, the size of Nck1 puncta grew and then shrank over time. This rise-and-fall pattern was common across puncta, regardless of the absolute lifetime of the individual punctum. Cross-correlation analysis of the intensity traces showed an average lag of approximately 5.2 seconds, with mNG-SHIP1 lagging slightly behind between the appearance of mCherry-Nck1 and subsequent arrival of mNG-SHIP1 (**Fig. 6D**). When we scored puncta based on arrival, peak, and departure using half-maximal intensity thresholds and peak timing, Nck1 preceded SHIP1 in more than 70% of co-localizing puncta for all three events (**Fig. 6E**). This temporal hierarchy is consistent with a model in which Nck1 helps nucleate puncta assembly at the membrane, while SHIP1 is subsequently recruited into the maturing complex via its C-terminal tail. Because Nck1 also departs before SHIP1 in most instances, Nck1 loss may represent the initial step in scaffold disassembly before SHIP1 dissociation.

**Figure 6.**
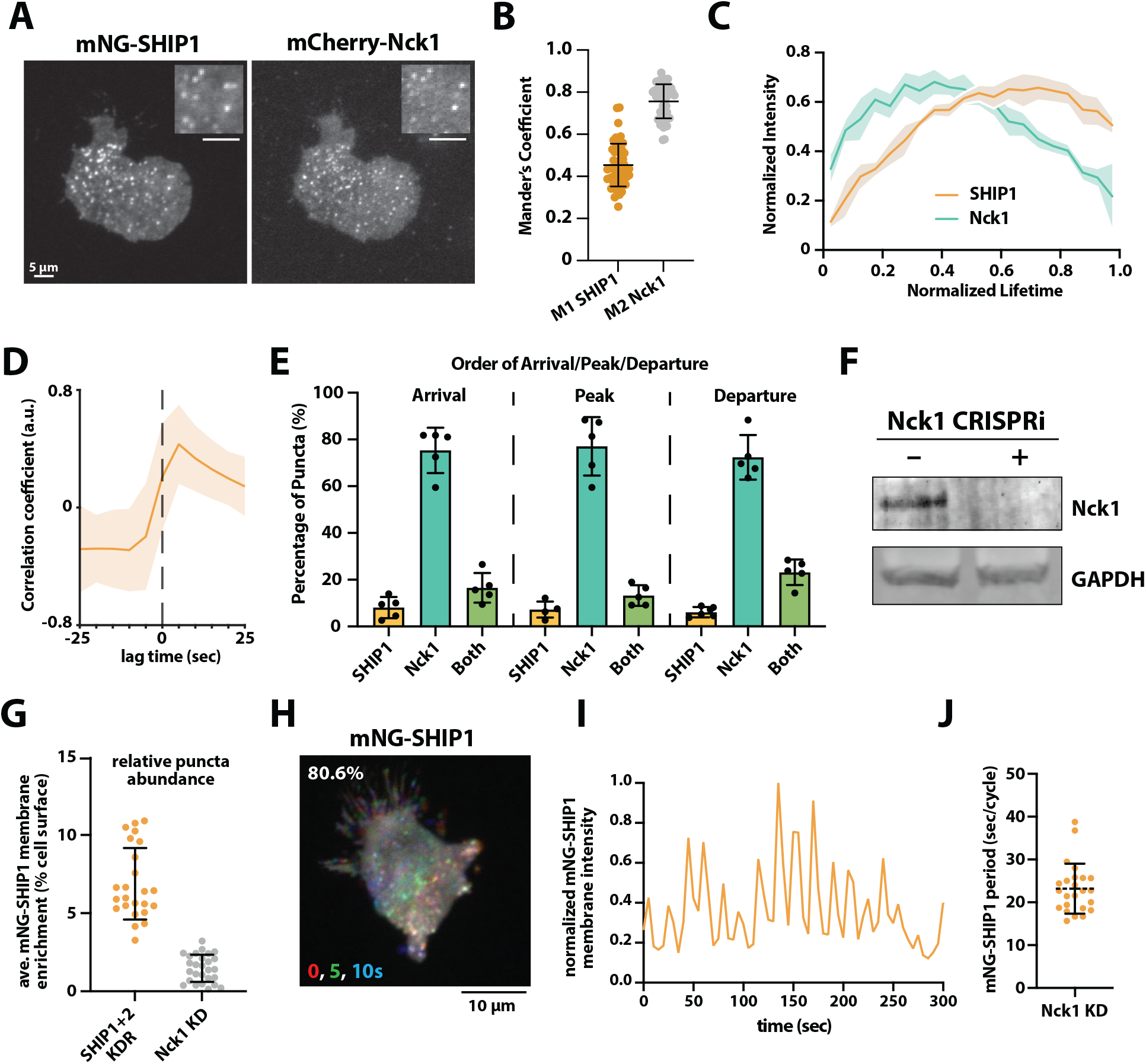
Nck1 regulates SHIP1 localization to membrane puncta. **(A)** Representative TIRF-M images showing localization of mNG-SHIP1 and mCherry-Nck1 in dHL-60. **(B)** Mander’s coefficients quantifying the fraction of SHIP1 overlapping with Nck1 (M1, 0.45 ± 0.1 by frames, orange) and vice versa (M2, 0.76 ± 0.1 by frames, gray). Points equal coefficient for a single frame (n = 18 total cells; N = 54 frames). **(C, D, E)** Nck1 temporally precedes SHIP1. **(C)** Normalized fluorescence intensity of co-localized SHIP1 (orange) and Nck1 (teal) puncta plotted against the normalized lifetime of individual puncta, where 0 represents appearance and 1 represents disappearance. Solid lines indicate the mean intensity calculated by averaging binned puncta lifetimes across independent cells (n = 5). **(D)** Representative mNG-SHIP1 and mCherry-Nck1 cross-correlation plot (6.0 ± 0.3 sec, n = 66 puncta). **(E)** Quantification of the order of arrival, peak, and departure for co-localized mNG-SHIP1 and mCherry-Nck1 puncta. Points equal average value per cells (n = 5). **(F)** Western blot showing expression of endogenous Nck1 in wild type and Nck1 CRISPRi dHL-60 cells. **(G)** Nck1 is required for mNG-SHIP1 membrane puncta localization. Quantification of exogenous mNG-SHIP1 membrane enrichment in SHIP1/2 CRISPRi control cells (n = 24, 6.88 ± 2.3 %) compared to Nck1 CRISPRi cells (n = 27, 1.48 ± 0.9%). Points equal mean enrichment per cell. **(H)** Representative temporal color-coded projection (RGB = 0, 5, 10s) of mNG-SHIP1 oscillating in a Nck1 KD cell. Phenotype observed in 80.6% of population (n = 36 cells). **(I)** Representative graph showing mNG-SHIP1 cortical membrane oscillations in Nck1 CRISPRi cells. **(J)** Autocorrelation-derived period length for mNG-SHIP1 oscillations observed in Nck1 CRISPRi cells (n = 24, 23.2 ± 5.8 sec). **(B**,**E**,**G**,**J)** Bars equal mean +/-SD. **(C-D)** Shaded bars equal SD.

To determine whether Nck1 is required for SHIP1 puncta localization, we silenced Nck1 expression using CRISPRi (**Fig. 6F**). While SHIP1 puncta normally occupied 6.9% of the total membrane surface area, we observed a 4.6-fold reduction (to 1.5%) in SHIP1 punctate localization in Nck1 knockdown cells (**Fig. 6G**). The fact that neutrophils also express Nck2 may explain the remaining SHIP1 localization to membrane puncta (Rincón et al. 2018). Interestingly, the decrease in mNG-SHIP1 localization to membrane puncta was correlated with an increase in mNG-SHIP1 localization to cortical oscillations (**Fig. 6H-6I**). We noticed that over time, dHL-60 cells expressing mNG-SHIP1 phenotypically stabilize into a more punctate-dominant localization mode, with cortical oscillations observed in around only 24.2% of cells. However, in Nck1 CRISPRi cells, 80.6% of mNG-SHIP1-expressing cells displayed oscillatory SHIP1 membrane dynamics, with a mean period of 23.2 seconds per cycle (**Fig. 6J**). This timing was similar to the cortical oscillations observed when mNG-SHIP1 was overexpressed in wild type cells (**Fig. 1E**), indicating that Nck1 depletion does not establish a new pathway for mNG-SHIP1 cortical oscillations. Together, these results indicate that SHIP1 partitions between cortical membrane oscillations and membrane puncta based on the relative expression of Nck1, FBP17, and SHIP1.

## DISCUSSION

Visualization of SHIP1 plasma membrane localization and dynamics in neutrophil-like cells using TIRF microscopy revealed localization to both leading-edge actin-based protrusions and cortical oscillations. When expressed at near-endogenous levels, however, SHIP1 localizes primarily to discrete membrane puncta with dynamics reminiscent of adhesion-receptor complexes. We discovered that the last 110 aa residues of the SHIP1 C-terminal domain are necessary and sufficient for plasma membrane localization to cortical oscillations and membrane puncta. While the SH2 domain has been shown to regulate SHIP1 localization in B-cell signaling, it was dispensable in neutrophils. The proline-rich motifs in SHIP1’s C-terminal tail provide a general mechanism for localizing SHIP1 to intracellular membranes through cell type-specific SH3 domain-containing protein interactions. Using CRISPRi knockdown-and-rescue experiments targeting SHIP1, we find that expression of a SHIP1 C-terminal domain truncation perturbs chemotaxis in a manner that phenocopies complete loss of SHIP1 function. In both cases, the neutrophils migrate slower and fail to directionally orient in the presence of a chemoattractant gradient.

In both resting and migrating neutrophils, ectopically expressed SHIP1 localizes to periodic traveling waves that propagate across the plasma membrane. Similar to the cortical oscillations observed in RBL-2H3 mast cells (D. Xiong et al. 2016), SHIP1 colocalizes with ectopically expressed FBP17 and a Cdc42(GTP) biosensor in neutrophils. Our findings indicate that overexpression of FBP17 and SHIP1 synergistically enhance cortical oscillations. This is likely caused by a shift in the steady-state distribution of interactions to favor FBP17-SHIP1 complex formation over other endogenous interactions. Nonetheless, membrane curvature sensing mediated by FBP17 is required for SHIP1 localization to the plasma membrane puncta and the leading edge of actin-based membrane protrusions. This was most evident when SHIP1 localization was visualized in dHL-60 cells. In this context, deletion of FBP17 and CIP4 abolished SHIP1 membrane puncta localization. By contrast, overexpression of FBP17 caused a redistribution of SHIP1 from predominantly membrane puncta to cortical membrane oscillations. This indicates that membrane curvature sensing proteins integrate upstream signals to initially establish SHIP1 membrane localization. A similar mechanism has been proposed for regulating SHIP2 localization to membranes enriched in endophilin and Lamellipodin (Chan Wah Hak et al. 2018).

The phosphatase activity of SHIP1 and SHIP2 has been implicated in regulating endocytosis (Boucrot et al. 2015). However, only the ubiquitously expressed SHIP2 has been shown to colocalize with endocytic markers (Nakatsu et al. 2010; Chan Wah Hak et al. 2018). Although the central phosphatase domain flanked by the PH and C2 domains is highly conserved between SHIP1 and SHIP2, the C-terminal domain sequences are divergent. In neutrophils, SHIP1 puncta dynamics resemble those of endocytic proteins, but they do not colocalize with clathrin or endophilin. This suggests SHIP1 plays a regulatory role, rather than a structural role, in endocytosis through the production of PI(3,4)P_2_ lipids. Recent work has shown that clathrin assembly drives the onset and direction of cortical waves through feedback from downstream endocytic factors including SHIP1, FBP17, Cdc42, and N-WASP (D. Xiong et al. 2016; Y. Yang et al. 2017). SHIP1 knockdown inhibits clathrin waves, implicating PI(3,4)P_2_ lipids as regulators of the frequency and the refractory period of cortical oscillations.

When we treat neutrophils with inhibitors that block substrate engagement of integrin receptors, we observe retrograde flow of SHIP1 membrane puncta coupled to the actin cytoskeleton. Although SHIP1 has been implicated in regulating adhesion-based cell migration in myeloid cells, we discovered that SHIP1 is part of a macromolecular assembly with localization dynamics reminiscent of adhesion receptors. Since prior studies relied on overexpression and used lower resolution imaging techniques, the SHIP1 membrane puncta localization has not been observed in the manner described here. Functioning downstream of membrane curvature sensing, we identified Nck1 as a key regulator of SHIP1 puncta formation. While proteins like FBP17 drive transient exploratory actin waves, Nck1 organizes the stable sites for SHIP1 localization. It remains unclear whether SHIP1 localization to cortical oscillations versus membrane puncta is beneficial for certain types of cell motility, such as adhesion-based versus amoeboid-based cell migration modes.

Nck1 and its paralog Nck2 are highly expressed in neutrophils (Rincón et al. 2018). Although Nck1 and Nck2 share 68% sequence identity and are partially redundant (Bladt et al., 2003), they have been shown to have distinct roles in functions including T cell receptor signaling, responses to growth factors, and regulation of the cytoskeleton (Borroto et al., 2014, Ngoenkam et al., 2014). These multivalent adaptor proteins consist of three SH3 domains and one SH2 domain (Buday et al., 2002). SHIP1 contains multiple proline-rich motifs (i.e. 969-974aa and 1040-1051aa) that can bind SH3-domain containing proteins. In addition, SHIP1 has multiple reported phosphotyrosine sites that serve as docking sites for SH2 domain containing proteins (Pauls and Marshall, 2017). In BCR-stimulated B cells, the SH2 domain of Nck1/2 has been shown to interact with pY685 and pY944 in the C-terminal domain of SHIP1 (Pauls et al., 2020). In neutrophils, we find that Nck1 localization consistently precedes SHIP1 recruitment to membrane puncta. When Nck1-dependent puncta are reduced by CRISPRi, SHIP1 localization shifts from predominantly puncta to cortical membrane oscillations that are presumably regulated by FBP17. These results suggest a dynamic higher-order assembly. In vitro and in vivo studies of nephrin, Nck, and N-WASP showed that Nck-containing multivalent networks can undergo phase-transition behavior and promote Arp2/3-directed actin assembly (Li et al., 2012). Biomolecular condensates often rely on this kind of multivalency to concentrate selected components and organize local reactions (Banani et al., 2017). Nck1 and SHIP1 colocalize in structures with nucleation and dissolution dynamics reminiscent of biomolecular condensates.

It remains unclear what biochemical state SHIP1 adopts when localized to cortical oscillations and membrane puncta. We previously found that the N-terminal SH2 domain regulates autoinhibitory interactions with the C2 domain that flanks the lipid phosphatase domain (Waddell et al. 2023; Drew et al. 2025). Although interactions with the CTD are required for SHIP1 membrane localization, it’s possible that a secondary interaction with tyrosine phosphorylated peptides may be required to fully activate SHIP1. Macromolecular molecular assemblies containing a mixture of SHIP1 and Nck1 could facilitate relief of autoinhibition. Alternatively, SHIP1-Nck1 membrane puncta could represent a sequestration mechanism meant to buffer the soluble concentration of SHIP1 and reduce either local or global phosphatase activity.

## DATA AVAILABILITY

All the information needed for interpretation of the data is presented in the manuscript. Plasmids related to this work are available upon request.

## ACKNOWLEDGMENTS

We thank Henry Rupp, Noah Smith, Meghan Chrissakis, and Cedric Stahel for assistance with molecular biology, cell culture, and live cell microscopy during the early stages of the project. We thank Peter Lechman (University of Oregon) for assistance cloning a CRISPR interference plasmid that expressed mNG.

## AUTHOR CONTRIBUTIONS

Resources: A.A., G.L.W., A.J.N., S.D., E.E.D., S.R.C. and S.D.H

Experiments and investigation: A.A., G.L.W., A.J.N., S.D., E.E.D., and S.D.H

Data Analysis: A.A., G.L.W., A.J.N., E.E.D., S.D., and S.D.H.

Conceptualization: A.A., G.L.W., and S.D.H.

Interpretation: A.A., G.L.W., and S.D.H.

Data curation: A.A., G.L.W., A.J.N., and S.D.H.

Writing – Review and editing: A.A., G.L.W., A.J.N., S.D., E.E.D., S.R.C. and S.D.H.

Writing – Original draft: A.A., G.L.W. and S.D.H.

Supervision: S.D.H.

Project administration: S.D.H.

Funding acquisition: S.D.H.

## FUNDING

Research was supported by the National Institute of General Medical Science R01 GM143406 (S.D.H.), NIH R01 GM143406 supplement (E.E.D.), R01 GM148769 (S.R.C.), and Molecular Biology and Biophysics Training Program T32 GM007759 (G.L.W. and E.E.D.). The content is solely the responsibility of the authors and does not necessarily represent the official views of the National Institutes of Health.

## CONFLICT OF INTEREST

The authors declare that they have no conflicts of interest with the contents of this article.

## MATERIALS & METHODS

### Molecular Biology

Genes encoding human INPP5D (SHIP1, 1-1189aa), FNBP1 (FBP17), Nck1 (1-377aa), clathrin light chain (CLTA, 1-248aa), endophilin A2 (SH3GL1, 1-368aa), endophilin B1 (SH3GLB1, 1-365aa), endophilin B2 (SH3GLB2, 1-395aa), and the CRIB domain (Cdc42(GTP) sensor) derived from the nucleation promoting factor, N-WASP, were PCR amplified with *Pfx* Accuprime mastermix in a T100 Bio-Rad thermal cycler. Following gel extraction and purification, DNA fragments were combined with the appropriate restriction enzyme digested plasmids and Gibson assembly was performed. Genes encoding the proteins described were expressed in frame fluorescent proteins in the following N-to C-terminal configuration: mNG-SHIP1, mCherry-SHIP1, EGFP-FBP17, TagBFP-FBP17, mCherry-Nck1, mCherry-CLTA, endoA2-mScarlet, endoB1-mScarlet, endoB2-mScarlet, and CRIB-iRFP. The complete open reading frame of all vectors used in this study were sequenced using Plasmidsaurus to ensure the plasmids lacked deleterious mutations. Refer to the Supplemental Text containing all the plasmid names.

### Cell culture

Differentiated neutrophil-like cells HL-60 (dHL-60s) are an established model system for studying cell polarity and migration of human leukocytes. These immortalized cells are derived from human patient with acute promyelocytic leukemia. In some studies, these cells are also referred to as PLB-985 cells, which is a subline of HL-60s. Single nucleotide polymorphism (SNP) analysis has found these cell line to be genetically identical (Rincón et al. 2018). Our cell line was originally obtained from the laboratory of Dr. Sean Collins (University of California at Davis) and is considered the PLB-985 subline of HL-60 cells. The exception is the HL-60 cell line containing genetic deletions of both FBP17 and CIP4, which we obtained from Orion Weiner (University of California at San Francisco). Cells were grown in suspension in RPMI 1640 + GlutaMAX media containing 25 mM HEPES (Life Technologies, cat #72400047), 9% fetal bovine serum (FBS), penicillin (100 units/ml) (Life Technologies, cat #15140122), streptomycin (100 µg/ml) (Life Technologies, cat #15140122). Cell lines were grown in humidified incubators at 37°C in the presence of 5% CO2 and split three times per week, keeping densities between 0.1-2 × 10^6^ cells/mL. The proliferating, undifferentiated HL-60 cells (uHL-60) were differentiate into a neutrophil-like cell type (dHL-60) by culturing 0.2 × 10^6^ cells/mL for 6 to 7 days in RPMI media supplemented with 2% FBS, penicillin (100 units/ml), streptomycin (100 µg/ml), 1.3% DMSO, and 2% Nutridoma-CS (Sigma, cat #11363743001). Nutridoma-CS was added to increase the chemotactic response of the cells, but this supplement is optional (Rincón et al. 2018). Human embryonic kidney (HEK) 293T Lenti-X were obtained from Takara (cat# 632180). These cells were transformed with adenovirus type 5 DNA and expresses the SV40 large T antigen. HEK293T Lenti-X cells were cultured in DMEM + GlutMAX + High Glucose (4.5 g/L) + sodium pyruvate (110 mg/L) (Life Technologies, cat #10569010) supplemented with 10% FBS (Sigma, cat# F4135-500ML), penicillin (100 units/ ml), and streptomycin (100 µg/ml). Cells were grown in 10 cm dishes in humidified incubators at 37°C in the presence of 5% CO_2_ and split at a confluency of 80-90% every 2-3 days. HEK293T Lenti-X cells were split using 1.5 mL of 0.25% Trypsin. Trypsin was quenched with 8.5 mL complete DMEM media containing 10% FBS. Cells were diluted 1:10 and seeded on a new 10 cm dish containing a total volume of 10 mL complete DMEM media warmed to 37°C.

### Live cell imaging

Extracellular Buffer (ECB: 5mM KCl, 125 mM NaCl, 1.5 mM CaCl_2_, 1.5 mM MgCl_2_, 10 mM glucose, 20 mM HEPES pH 7.4) and Leibovitz-15 (Life Technologies, catalog #21083027) complete media (10% FBS, 100 U/mL Pen/Strep) were warmed to 37°C and combined 1:4 to make the Imaging Media. Differentiated PLB-985 cells were prepared for imaging by centrifuging 0.5 mL of cells at 100 xg for 10 minutes, aspirating off the medium, and resuspending the cells in warm imaging media (L-15 complete media to ECB, 1:4 parts). Differentiated PLB-985 cells were then imaged using one of two different live cell imaging methods: a method using fibronectin coated glass attached to an IBIDI flow cell chamber or a previously described 96-well format under-agarose method (Bell et al. 2018). To image cells on fibronectin coated glass, 25 × 75 mm coverslips were cleaned with 2% Hellmanex III (ThermoFisher, Cat#14-385-864), washed with MilliQ water, dried with N_2_ gas, and attached to an IBIDI flow cell chamber (IBIDI sticky-Slide VI 0.4, catalog #80608). A 10 µg/mL fibronectin (Sigma, cat # F1141, 1 mg/mL stock concentration) solution diluted in 1x PBS was added to each well of the IBIDI chamber, incubated for 30-60 minutes, and unbound fibronectin was washed out with 1x PBS. 100 µL of differentiated PLB-985 cells were flowed into the IBIDI chamber. Cells would adhere within 5-10 minutes. Cells were imaged in the presence of uniform chemoattract by flowing in 100 µL of 20 nM fMLF. To image cells under agarose, 5 µL of differentiated PLB-985 cells were spotted in the center of a well in a glass-bottom 96 well plate (cat #) and allowed to settle by gravity onto the bottom of the glass for 5 minutes. A 3% wt/vol low melting agarose (Gold-Bio, Cat # 204-100) solution (3% LMA) was prepared in ECB, heated to 70°C to ensure the agarose was fully dissolved, and combined 1:1 with warmed imaging media so that the final concentration of the LMA is 1.5% wt/ vol. 195 µL of this warm 1.5% LMA mixture is then gently pipetted above the 5 µL of differentiated PLB-985 cells. Slowly pipetting down the side of the well is the best way to avoid disturbing the 5 µL of settled cells. The plate was covered with aluminum foil to protect it from light and left at room temperature for 10 minutes to allow the agarose to solidify. After 10 minutes the cells could be immediately imaged or allowed to incubate for another 10 minutes in a 37°C incubator before starting the imaging experiment. This method can also be used to image cells in the presence of a chemoattractant gradient. We prepared 2x solutions of caged fluorescein (CMNB-F (2x = 2 µM), Life Technologies, Cat# F-7103) or caged chemoattractant *N*-nitroveratryl fMLF (nv-fMLF) (2x = 300 nM) in imaging media as previously described (Collins et al. 2015; Pirrung et al. 2000). These solutions were combined 1:1 with the warm 3% LMA mixture and gently plated above the cells as described above. During an imaging experiment, a UV laser can be used to uncage nv-fMLF to create a chemoattractant gradient.

### Lentivirus production

Lentivirus was generated by transfecting 60-70% confluent HEK293 Lenti-X cells in a 10-cm plate containing 8 mL of complete media. Transfection reagents were prepared by combining 6.7 µg psPAX2 (2nd generation lentiviral packaging plasmid, Addgene #12260), 0.85 µg pVSV-G (Expresses VSV-G envelop protein for pseudotyping NanoMEDIC particle, Addgene #138479) (Gee et al. 2020), 7.5 µg of transfer lentiviral vector containing gene of interest, and 30 µL of 1 mg/mL polyethyleneimine (Polyscience, Cat#) in 0.5 mL Opti-Mem (ThermoFisher, cat#31985070). This mixture was incubated for 15 minutes at room temperature before adding dropwise to plated HEK293 Lenti-X cells. Media containing lentivirus was harvested 48 hours after initiating transfection and clarified by centrifugation to remove cell debris. Lentivirus was concentrated by adding 333 µL Lenti-X concentrator (Takara, cat# 631231) per mL of viral supernatant, incubated overnight at 4°C, and then centrifuged at 1500 × g for 45 minutes. Supernatant was aspirated and the white viral containing pellet was resuspended in 0.4 mL of complete RPMI media (9% FBS). Lentivirus was used immediately or stored at -80°C. To infect PLB-985 cells, 0.4 mL of concentrated lentivirus was added with 5 mL of undifferentiated cells at a density of 0.2 × 10^6^ cells/mL containing a final concentration of 8 µg/mL polybrene (Millipore, Cat# TR-1003-G) in the cell culture media. The cells were passaged at least one time before differentiating into the neutrophil-like state (see ***Cell Culture*** for protocol).

### CRISPR interference

For CRISPRi, 5 mL of undifferentiated PLB-985 cells at a density of 0.2 × 10^6^ cells/mL were transduced by adding pHR-UCOE-EF1a-KRAB-dCas9-P2A-BLASTICIDIN (Addgene, Cat#85969) lentivirus in the cell culture media containing 8 µg/mL polybrene (Millipore, Cat# TR-1003-G) (see ***Lentiviral production*** for protocol). This construct contained a minimal ubiquitous chromatin opening element (UCOE) was inserted upstream of the SFFV promoter. After 3 days of infection, selection of the dCas9-KRAB construct was initiated by adding 10 µg/ mL blasticidin. After 7 days of antibiotic selection the dCas9-KRAB expressing PLB-985 cells reached a growth rate that mirrored the non-transduced cells. To silence SHIP1 and Nck1, dCas9-KRAB expressing cells were transduced with mU6-sgRNA-SHIP1-PURO and pU6-gRNA-Nck1-EF1a-PURO-T2A-TagBFP respectively. Guide RNA designs were based Benchling [Biology software] and the Dolcetto library (Sanson et al. 2018). Guide RNA protospacer sequences were individually cloned into both the sgRNA expression vector using the BstXI and BlpI restriction sites and T4 DNA ligase-based cloning. Each plasmid was then verified by Sanger sequencing of the protospacer. For genetic rescues, the plasmid containing the sgRNA sequence was modified to co-express the desired rescue construct on a single vector. This maintains consistent expression stoichiometry and prevents the variability associated with triple-plasmid transductions. Several SHIP1 rescue variants were generated, including full-length mNG-SHIP1(1-1189aa) and the C-terminal truncation mutant (i.e. 1-1078aa).

### Immunoprecipitation

PLB-985 cells (at least 10 million per sample) were resuspended in 500 μL Solubilization Buffer (50 mM Tris pH 7.5, 5 mM EDTA, 150 mM NaCl, 1% Triton X-100) and 10 μL Protease Inhibitor Cocktail (Sigma-Aldrich, Cat. # P8340) and rotated for 1 hour at 4°C. A sample of whole cell lysate was removed, and total protein was quantified using the Pierce BCA Protein Assay Kit (Thermo Fisher, Cat # 23225). A second whole cell lysate sample was collected and mixed 4:1 with 4X SDS-PAGE sample buffer supplemented with 2% β-mercaptoethanol (BME). 50 μL Protein A Dynabeads (invitrogen, Cat. # 10001D) were washed with 500 μL PBS-T (1xPBS with 0.1% Tween-20) and resuspended in a solution of 150 μL PBS-T and 50 μL appropriate primary antibody (Santa Cruz, Cat. # 8425, 166641, 515414). After rotating for 10 minutes, beads were pelleted and resuspended in 200 μL PBS-T. Beads were pelleted, resuspended in the cell lysate (total protein amount was normalized between samples using values obtained from the BCA Protein Assay), and rotated for 4-6 hours at 4°C. Beads were pelleted and resuspended in 40 μL 1X SDS-PAGE sample buffer supplemented with 2% β-mercaptoethanol and boiled for 15 minutes. Beads were pelleted and supernatant (enriched lysate) was collected.

### Western blot

Enriched and whole cell lysate samples were loaded on Mini-PROTEAN TGX precast gels (Bio-Rad, Cat. # 4561094) along with the Precision Plus Protein Dual Color Ladder (Bio-Rad, Cat. # 1610374). Protein bands were separated at 200V for 35 minutes. Gels were transferred to nitrocellulose membranes (Bio-Rad, Cat. # 1704271) using cold Transfer Buffer (40ml 5x Transfer Buffer (Bio-Rad, Cat. # 1704271), 40ml ethanol, 120ml miliQ water) and the 7-minute mixed MW protocol on the Bio-Rad Trans-Blot Turbo Transfer System (Bio-Rad, Cat. # 1704150). Blots were washed with MiliQ water and stained with 0.1% Ponceau in 5% acetic acid for 5 minutes. Blots were washed with MiliQ water and blocked for at least 1 hour with Intercept Blocking Buffer (LI-COR, Cat. # 927-70001). Blots were incubated at 4°C overnight in 1:250 appropriate fluorescent antibody (Santa Cruz, Cat. # 8425 Ax647, 166641 Ax488, 515414 Ax546) and 1:500 mouse anti-GADPH primary antibody (Santa Cruz, Cat. # 32233) in Blocking Buffer. Blots were washed four times for 5 minutes with PBS-T and scanned on the Amersham Typhoon (Cytiva). Blots were incubated at 4°C overnight in 1:1000 anti-mouse IgGκ (Santa Cruz, Cat. # 516180) in Blocking Buffer, washed, and scanned.

### Microscope hardware and imaging acquisition

Live cell TIRF microscopy experiments were performed using an inverted Nikon Ti2 microscope. All images were acquired using an iXon Life 897 EMCCD camera (Andor Technology Ltd., UK). A Nikon 100x oil immersion objective (1.49 NA) was used for all single molecule TIRF experiment. Cortical oscillations were visualized in PLB-985 cells using a Nikon 60x oil immersion objective (1.49 NA). Fluorescently labeled proteins were excited with either a 405, 488, 561, or 647 nm diode lasers (Coherent Inc. Santa Clara, CA) controlled with a Vortran (Sacramento, CA) laser driver digital modulation. The following the laser powers measured through the objective were used to excite mNG-SHIP1 (1 mW 488 nm), iRFP-CRIB (1 mW 647 nm), mCherry-SHIP1 (1 mW 561 nm). Excitation light was passed through the following dichroic filter cubes before illuminating the sample: (1) ZT488/647rpc and (2) ZT561rdc (ET575LP) (Semrock). Nikon emission filter wheel housing 25mm ET525/50M, ET600/50M, and ET700/75M filters (Semrock). Unless otherwise noted, live cell imaging experiments were performed at room temperature (i.e. 23^º^C). All microscope hardware was controlled with NIS elements. Live cell chemotaxis assays involving nv-fMLF were performed under brightfield microscopy. These experiments utilized a Nikon 20x (0.4 NA) objective, which provided a wider field of view suitable for capturing long-range cell trajectories. Time-lapse sequences were acquired at 37C using a heated stage, and images were taken at intervals of 2s. After imaging baseline conditions for 10 minutes, an initial UV uncageing was performed for 5s. Subsequent uncageings were performed for 20 ms at intervals of 1.5 minutes.

### Image processing and data analysis

Image analysis was performed on ImageJ and MATLAB Curve fitting was performed using MATLAB and Prism 9 (GraphPad). The power spectrum, autocorrelation, and cross-correlation graphs were analyzed through an application created through MATLAB App Designer. The input is an Excel file upload with x and y values of time and fluorescence intensity respectively.

### Power Spectrum

Generated by processing the fluorescence signal using MATLAB’s built-in fast Fourier transform (FFT) algorithm and multiplying the processed sequence by its complex conjugate.

Y = fft(X)

Zc = conj(Z)

MATLAB’s FFT algorithm is based on the discrete Fourier transform (DFT). Under the assumption that the sig nal is periodic, The DFT breaks down the input signal in the following way, where X, N, n, and k are the transformed sequence, number of samples, time-domain index, and frequency-domain index respectively:

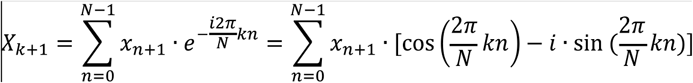

The resulting output is a complex number that is described by an “imaginary” sine component and a “real” cosine component. Inputting a signal consisting of solely cosines would return only real values, and inputting a signal consisting of solely sines would return only imaginary values. Any fluorescence signal, however, would require both components as the imaginary component is necessary to represent its phase shifts. As such, while the power spectrum of a signal can normally be calculated simply by squaring the magnitude of each frequency, the imaginary component to the DFT equation poses an issue as the following error would appear: “Imaginary parts of complex X and/or Y arguments ignored”. To account for the imaginary component, the complex conjugate of the transformed signal was taken instead.

### Autocorrelation analysis

The autocorrelation analysis was conducted using MATLAB’s autocorrelation function, [acf,lags] = autocorr(y). The resulting correlation coefficients were plotted against phase lag.

### Cross-correlation analysis

The cross-correlation analysis was conducted using MATLAB’s cross-correlation function, r = xcorr(x,y). The resulting correlation coefficients were plotted against phase lag.

The input requires uploads of two separate Excel data files, each with time and fluorescence intensity as x and y values respectively.

### DFT derivations

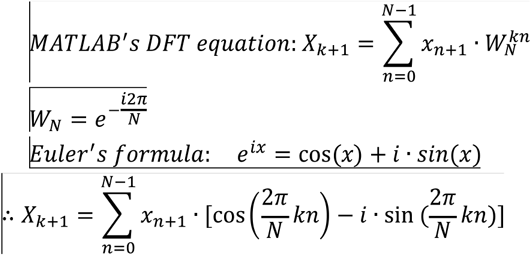

### Quantification of cell morphology

Binary masks were generated for each cell using custom MATLAB scripts. For brightfield images, segmentation was performed using the Segment Anything Model (SAM) via a MATLAB interface (Kirillov et al. 2023). For fluorescence images, masks were generated using a MATLAB application that enables edge detection using a Sobel operator with a user-defined scaling factor for thresholding, dilation of detected edges using linear structural elements, closing of internal gaps within the dilated boundaries, and user-defined size filtering. For each identified cell, a series of morphological features was extracted using the region props function and other geometric calculations. For multi-parametric analysis, 2D scatter plots were generated with integrated marginal Kernel Density Estimation (KDE) plots, allowing for the visualization of both individual cell correlations and the overall density distribution of the population across selected metrics.

### Quantification of neutrophil chemotaxis

#### Segmentation and centroid-tracking

Initial cell segmentation was performed using Cellpose (Stringer et al. 2021), a deep learning-based cellular segmentation algorithm. Custom models were trained using a representative subset of frames from each experimental condition. The resulting segmentation masks were used to define the boundaries of every cell in each frame of the video. Following segmentation, the binary masks were imported into MATLAB, where the geometric centroid of each identified cell was calculated. These centroid coordinates served as the standardized input for the subsequent tracking algorithms.

#### Trajectory reconstruction

Cell positions were linked between consecutive frames using a nearest-neighbor distance algorithm with a user-defined maximum linking displacement threshold. Only cells tracked for a minimum of 10 consecutive frames were indexed and retained for kinematic analysis.

#### Kinematic analysis

Each individual track was first translated so that its initial coordinates at its starting time point (t0) were set to the origin (0,0) of a Cartesian plane. The precise spatial coordinate of the chemoattractant source was determined empirically for each experiment using an nv-fluorescein control; by visualizing the area of UV-induced fluorescence, the center of the illuminated region was defined as the chemoattractant source. Using this source coordinate, a rotation matrix was applied to the trajectory to align the vector between the cell’s initial position and the source with the positive y-axis (i.e this vector is the idealized cell trajectory). Individual trajectories were gradient-colored by their respective durations (blue at t_0_ to red at t^final^). This spatiotemporal normalization enables a visual comparison of chemotactic efficiency across experimental conditions. To provide a spatial reference for the chemoattractant source, a black triangle was plotted along the positive y-axis. The distance of this marker from the origin was defined as the median distance from the initial starting position to the chemoattractant source across all tracks in the dataset. Following normalization, chemotactic behavior was quantified on a per-vector, per-step basis. The cell speed for each step was calculated as the magnitude of the displacement vector divided by the time interval. Angular bias was determined by calculating the angle between each stepwise movement vector and the idealized source vector (i.e the +y vector), then subtracting that number from 90º such that an angular bias of 90º and 0º represent perfect chemotaxis and random movement respectively. Directed speed was calculated as the scalar component of the total velocity that contributes specifically to displacement along the idealized source vector. All population-level metrics were calculated using displacement–weighted averages. For example, if a cell takes a 10 µm step toward the source at a 90° bias followed by a 1 um step at a 0° bias, the resulting weighted average would be 81.8°. In contrast, a simple average would over-represent the noise and drop the score to 45°.

### Quantification of puncta dynamics

#### Puncta probability distribution and spatial mapping

Puncta were identified using a Difference of Gaussians (DoG) filter, binarized using a user-tuned intensity threshold, and linked across frames using a nearest-neighbor tracking algorithm. To map puncta coordinates, the cell’s polarity axis was defined using a hybrid approach based on the cell’s motility state. (1) For migrating cells, orientation was determined using a smoothed look-ahead velocity vector, where centroid positions were smoothed using a 10-frame Gaussian window, and the orientation vector was calculated as the vector between the current smoothed centroid and the centroid 5 frames ahead. (2) In experimental conditions where cells remained stationary but polarized (e.g EDTA treatment), orientation was derived from the major axis of an ellipse fit to the binary mask and a one-time user selection of the leading-edge point. To account for variations in cell length and morphology, puncta positions were mapped onto a normalized longitudinal scale. For each frame, the cell mask was rotated to align the leading edge to the positive y-axis and sliced into 1000 horizontal bins. The y-coordinates for the puncta were rescaled to range of [0, 1], where 1 represents the leading edge and 0 represents the trailing edge. An average cell shape was reconstructed by averaging the width profiles across all frames for each bin. Spatial probability density functions were calculated to visualize, relative to the leading edge, the organization of puncta at their points of appearance and disappearance. The puncta densities were smoothed using a Gaussian kernel and plotted as heatmaps overlaid onto the average cell shape. Net flux was mapped by subtracting the disappearance probability from the appearance probability.

#### Temporal dynamics of co-localized puncta

To determine the sequential order of arrival, peak intensity, and departure for co-localized puncta across two channels, candidate co-localization between channel 1 (Ch1) track *i* and channel 2 (Ch2) track *j* was first established using a frame-by-frame spatial voting system. For each frame where track *i* was present, the nearest Ch2 track within a user-defined distance threshold receives a single vote. The Ch2 track accumulating the highest number of votes was designated as the candidate partner. To be accepted as a validated co-localized pair, the total frame overlap count was required to satisfy a fractional duration threshold:

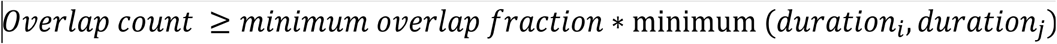

Following track validation, the masks belonging to the structurally better-resolved channel was used as a spatial reference to monitor fluorescence intensity over time across both channels. To account for varying event durations and enable population-level comparisons, puncta intensities were normalized, and puncta lifetimes were rescaled into 20 equidistant temporal bins via linear interpolation. The relative timing of puncta dynamics (i.e., arrival, peak, and departure) was calculated and compared between channels using maximum and half-maximum intensity thresholds.

#### Cross-correlation analysis

To determine the temporal offset (i.e lag time) between paired puncta, per-pair cross-correlation analysis was conducted using mean-subtracted, normalized intensity time series using MATLAB’s xcorr function in coeff mode. To eliminate overfitting on short trajectories, the maximum allowable lag was constrained to 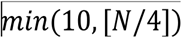 frames, where 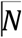 is the number of valid, non-NaN frames in the pair. Discrete peak lags were refined to sub-frame precision using parabolic interpolation across the three data points flanking the discrete maximum.

## Notes

### Competing Interest Statement

The authors have declared no competing interest.

